# Antibiotic treatment modulates *Escherichia coli*-derived bacterial extracellular vesicle (BEV) production and their capacity to upregulate ICAM-1 in human endothelial cells

**DOI:** 10.1101/2025.05.12.653517

**Authors:** Louis P. Widom, Panteha Torabian, Abigail C. Wojehowski, Sina Ghaemmaghami, Lea V. Michel, Thomas R. Gaborski

## Abstract

Antibiotic treatment is often necessary to eliminate life-threatening bacterial infections. However, these treatments can alter production of bacterial extracellular vesicles (BEVs), which often contain pro-inflammatory biomolecules. In this study, we examined how the clinically-relevant antibiotics meropenem, tobramycin, and ciprofloxacin impacted BEV production from a urinary tract infection-associated *Escherichia coli* strain (CFT073 [WAM2267]) and a meningitis-associated strain (K1 RS218). BEVs from both strains caused a dose-dependent increase in expression of intercellular adhesion molecule-1 (ICAM-1) in human umbilical vein endothelial cells, priming the endothelium for interactions with immune cells. Blockade of toll-like receptor 4 revealed that this receptor was responsible for BEV-endothelial interactions. Treatment with meropenem, a β-lactam antibiotic, increased production of BEVs from strain K1 RS218. Furthermore, meropenem treatment caused strain CFT073 [WAM2267] to produce BEVs with heightened stimulatory capacity, possibly by amplifying the content of lipoprotein Lpp in these BEVs as measured by mass spectrometry. To our knowledge, this is the first study examining the interplay between antibiotic treatment and the effects of the resulting BEVs on endothelial ICAM-1 expression. Our results indicate treatment risks of certain antibiotics against specific strains of *E. coli* and could help identify therapeutic targets to reduce BEV-mediated endothelial stimulation during infection.

## Introduction

Sepsis arises from an exaggerated immune response to infection, leading to widespread inflammation and tissue damage^1^. It remains a global health crisis, with approximately 48.9 million cases and 11 million deaths reported annually, underscoring the urgent need for effective prevention and treatment strategies^2^. A critical component of this inflammatory response is intercellular adhesion molecule-1 (ICAM-1), a 90 kDa member of the immunoglobulin superfamily whose expression is upregulated in response to pro-inflammatory cytokines^3^. ICAM-1 is essential for leukocyte arrest and transmigration and is constitutively present on endothelial cells^3^. In sepsis, these processes can lead to immune cell infiltration into the brain, exacerbating disease outcomes by increasing the likelihood of developing cognitive impairment^4–6^. ICAM-1 upregulation in the endothelium may also occur due to exposure to bacterial products such as lipopolysaccharide (LPS), which is a major outer membrane component of gram-negative bacteria^7^. In other words, bacteria and their secretions can prime the endothelium for interactions with leukocytes that can contribute to sepsis-related inflammatory pathologies.

Bacterial extracellular vesicles (BEVs) are small, 40–400 nm lipid-membrane-bound particles that are secreted by both gram-negative and gram-positive bacteria^8^. Their cargo includes various nucleic acids, signaling factors, and toxins^9^. In gram-negative bacteria such as *Escherichia coli* (*E. coli*), many of these vesicles are derived from the outer membrane and are alternatively referred to as outer membrane vesicles^10^. As such, they can contain membrane components such as LPS and outer membrane proteins, and these factors can promote an inflammatory response^11,12^. One interesting possibility is that blood-borne BEVs may persist longer in the circulation compared to smaller free bacterial factors. For example, the half-life of free LPS in mice is only 2–4 min whereas mammalian extracellular vesicles have a half-life of 30–360 min in blood^13,14^. The stability of *E. coli*-derived BEVs may also be aided by expression of anti-phagocytic proteins that could inhibit clearance by immune cells, so there is potential for BEVs to travel longer, penetrate deeper into tissues, and thus have more opportunities to induce both acute and sustained pro-inflammatory stimulation than smaller bacterial factors^15,16^.

Although antibiotics are critical for infection control, they can paradoxically enhance BEV release, which may deliver pathogenic factors that alter host cell responses^17–19^. Investigating the role of these antibiotic-induced BEVs in driving ICAM-1 upregulation is vital, as this process can exacerbate inflammation and contribute to endothelial dysfunction, a hallmark of sepsis pathogenesis^3^. Notably, prior research has demonstrated that BEVs can significantly elevate endothelial ICAM-1 expression^20^. However, to our knowledge, it has not yet been determined how antibiotic treatment modulates the ability of BEVs to activate endothelial cells and upregulate ICAM-1 expression.

We investigated two *E. coli* strains to determine if the effects of antibiotic treatment on BEV production could vary in a strain-dependent manner. The first was uropathogenic *E. coli* strain CFT073 [WAM2267], responsible for causing urinary tract infections (UTIs), which are the second most common infectious diseases in humans after respiratory infections^21^. UTIs occur when pathogenic bacteria, predominantly *E. coli*, invade urothelial cells, triggering inflammatory responses in the urethra that result in symptoms such as dysuria, perineal discomfort, and increased urinary frequency and urgency^22^. The CFT073 strain is particularly virulent, capable of colonizing the bladder and ascending to the kidneys, which can lead to conditions such as urosepsis. Uropathogenic *E. coli* employ various pathogenic mechanisms including fimbrial adherence, toxin production, and evasion of host defenses^23^. The second strain, K1 RS218, was isolated from the cerebrospinal fluid of a neonate with meningitis. This strain is particularly significant due to its ability to cross the blood-brain barrier, a critical step in causing neonatal meningitis. K1 RS218 achieves this through specialized virulence factors, such as the K1 capsule, which provides resistance to host immune defenses, facilitating the development of severe inflammation and neurological damage in newborns^24,25^. K1 *E. coli* can also initiate sepsis^26^. Like other gram-negative bacteria, both of these *E. coli* strains produce LPS, which can induce pro-inflammatory signaling by binding to the extracellular domain of toll-like receptor-4 (TLR4) expressed on endothelial cells, immune cells, and various other cell types^27^. Multiple TLR4 inhibitors have been investigated as a means to improve outcomes from severe sepsis, but so far these approaches have been unsuccessful in clinical trials^28,29^. This may be due to the complexity of sepsis and/or the involvement of various other bacterial products during infection.

To examine the impact of antibiotics on *E. coli* BEV production and stimulatory capacity, we selected meropenem, tobramycin, and ciprofloxacin because of their varying modes of action as well as their clinical relevance and efficacy against UTIs, urosepsis, and bacterial meningitis^30–34^. Meropenem is a broad-spectrum β-lactam antibiotic effective against severe infections. It exerts its bactericidal action by binding to penicillin-binding proteins in the bacterial cell wall, inhibiting peptidoglycan crosslinking and cell wall synthesis, which ultimately leads to cell death. Its utility in treating resistant infections is emphasized in clinical guidelines^35^. Tobramycin, an aminoglycoside antibiotic, is particularly effective against gram-negative bacteria, including *E. coli*, making it a valuable treatment option for complicated infections^33^. In clinical studies, 21 out of 30 patients with severe or complicated gram-negative UTIs were cured after a 5-day course of tobramycin, with no reported side effects^36^. It has also been used to successfully treat child and adult bacterial meningitis^33^. Ciprofloxacin, a third-generation fluoroquinolone, is a key treatment for UTIs, particularly those caused by *E. coli*^22^. Its effectiveness is due to its ability to inhibit bacterial DNA gyrase, preventing DNA synthesis and arresting bacterial growth. Additionally, ciprofloxacin’s high drug permeability allows it to quickly reach therapeutic concentrations in prostatic fluid and bladder urine, resulting in a rapid bactericidal effect^22^.

The present study sought to understand whether these three different classes of antibiotics would have varying effects on *E. coli* BEV production and the capacity of the secreted BEVs to promote endothelial activation. These interactions were studied with both CFT073 [WAM2267] and K1 RS218 to determine whether there might be strain-specific differences in *E. coli* BEV release in response to antibiotic treatment. We found that there were indeed antibiotic- and strain-dependent differences in the BEVs produced by these *E. coli* that resulted in altered effects on endothelial expression of ICAM-1. The results from this study may help inform clinicians about the risks of treating infected patients with certain antibiotics depending on which bacterial strains are present.

## Results

### BEVs cause increased endothelial ICAM-1 expression in a dose-dependent manner

To investigate the potential of BEVs to promote endothelial activation, we examined expression of inflammatory marker ICAM-1 on the surface of human umbilical vein endothelial cells (HUVECs) following 16–17 h exposure to BEVs derived from two *E. coli* strains: CFT073 [WAM2267] and K1 RS218. Immunofluorescence staining showed a dose-dependent increase in ICAM-1 expression in response to BEVs from both strains, with significant increases above baseline at concentrations of 1E6 BEVs/ml (approximately 3.30 BEVs/HUVEC) for the CFT073 strain and at 1E7 BEVs/ml (approximately 37.8 BEVs/HUVEC) for the RS218 strain (Fig. 1). This indicated that the CFT073-derived BEVs were more potent than the RS218-derived BEVs, despite the fact that they were obtained from two strains of the same bacterial species. It is therefore possible that strain-specific variations exist in the composition of BEVs that can influence their stimulatory capacity. Additionally, the immunofluorescence images indicated some heterogeneous upregulation of ICAM-1 even at lower BEV concentrations, which could suggest HUVEC sensitivity to individual BEVs. For both strains, ICAM-1 upregulation following BEV exposure was comparable to that induced by 10 ng/mL LPS, which was used as a positive control for increased ICAM-1 expression. This amount of LPS is over twenty times the concentration that has been observed in patient serum during sepsis, and should therefore elicit a consistently high ICAM-1 response^37^. The results from these dosing experiments support the idea that BEVs play an active role in modulating endothelial cell function and could contribute to vascular inflammation in the context of bacterial infections. Because treatment with 1E8 BEVs/ml from both *E. coli* strains routinely promoted high ICAM-1 expression by HUVECs, this concentration was selected for all of the following cell stimulation experiments except where otherwise indicated. Note that this concentration corresponded to approximately 322.6–385.4 BEVs/HUVEC in these dosing experiments, but we have decided to express concentrations in terms of BEVs/ml throughout this manuscript to avoid the assumption that 100% of the BEVs settled to the bottom of the solution to interact with the HUVEC monolayer during the treatment period.

**Fig. 1:**
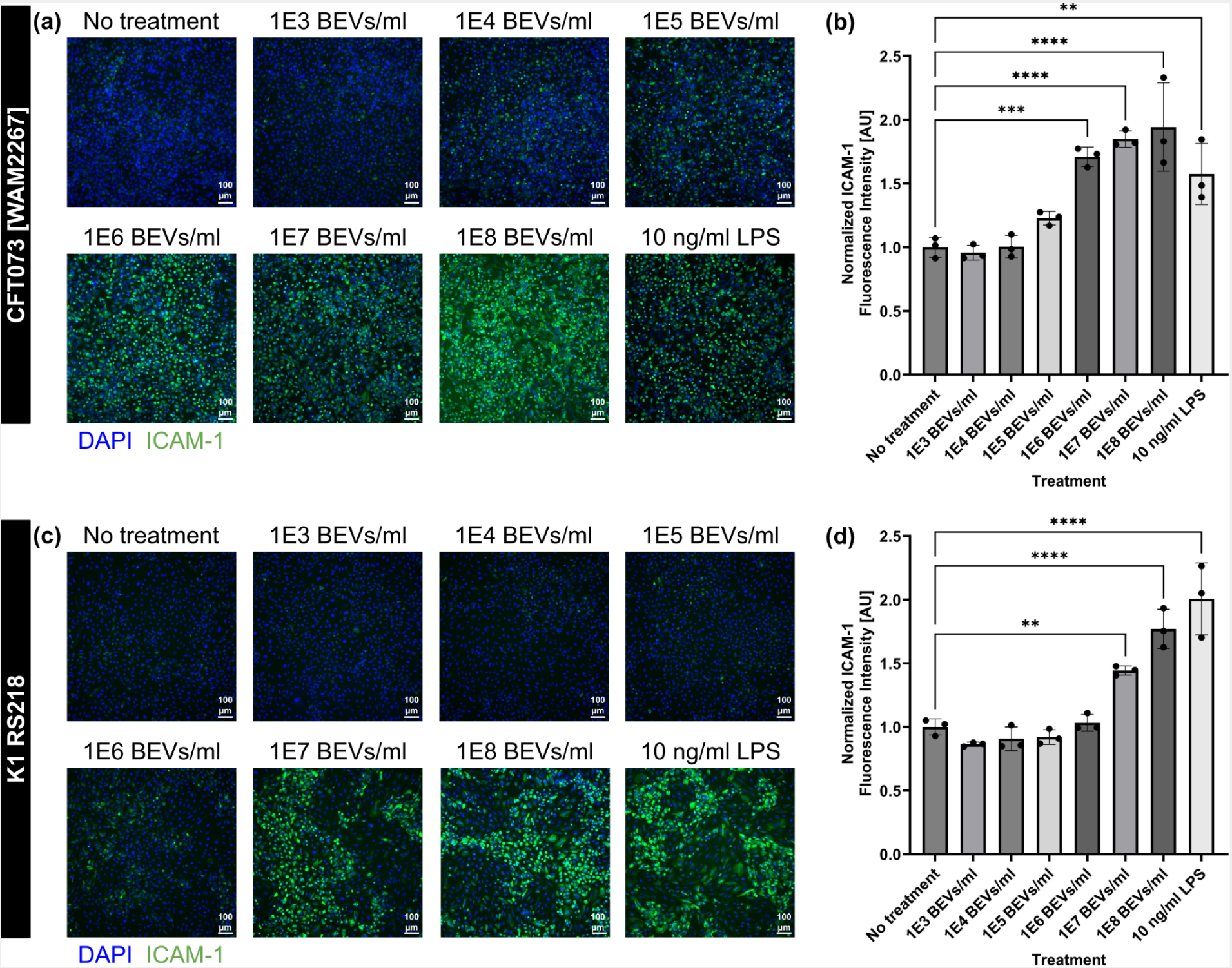
HUVEC ICAM-1 expression increases due to exposure to *E. coli*-derived BEVs in a dose-dependent manner. (**a**) Immunofluorescence images of HUVECs stained with DAPI (blue) and anti-ICAM-1 antibody (green). Cells were treated with increasing concentrations of BEVs derived from *E. coli* strain CFT073 [WAM2276] or with 10 ng/ml LPS as a positive control for ICAM-1 expression for 16–17 h. (**b**) Barplot of ICAM-1 fluorescence intensity from HUVECs treated with BEVs derived from *E. coli* strain CFT073 [WAM2276]. The average fluorescence intensity of ICAM-1 in each image was divided by the number of cell nuclei in the corresponding DAPI channel. These values were then normalized to the average for the wells that received no stimulatory treatment. (**c**) Immunofluorescence images of HUVECs stained with DAPI (blue) and anti-ICAM-1 antibody (green). Cells were treated with increasing concentrations of BEVs derived from *E. coli* strain K1 RS218 or with 10 ng/ml LPS as a positive control for ICAM-1 expression for 16–17 h. (**d**) Barplot of ICAM-1 fluorescence intensity from HUVECs treated with BEVs derived from *E. coli* strain K1 RS218. The average fluorescence intensity of ICAM-1 in each image was divided by the number of cell nuclei in the corresponding DAPI channel. These values were then normalized to the average for the wells that received no stimulatory treatment. Error bars = s.d. Performed one-way ANOVA with Dunnett post-hoc test: ✱✱ = *p* < 0.01, ✱✱✱ = *p* < 0.001, ✱✱✱✱ = *p* < 0.0001

### *E. coli*-derived BEVs cause a stimulatory response by binding to TLR4 on HUVECs

To investigate whether LPS was a primary stimulatory component of these *E. coli*-derived BEVs, we blocked TLR4 by treating HUVECs with inhibitor TAK-242 (CAS 243984-11-4). Since TLR4 is the receptor responsible for binding and sensing LPS, this inhibition would hypothetically prevent BEVs from causing increased ICAM-1 expression in HUVECs. First, we confirmed TAK-242’s specificity for TLR4 by stimulating HUVECs with either LPS or with a pro-inflammatory cytomix consisting of TNF-α, IL-1β, and IFN-γ. The cytomix components are known to bind to the receptors TNFR1, TNFR2, IL-1R1, and IFNGR rather than TLR4^38–40^. When we treated HUVECs with LPS and TAK-242 there was no increase in ICAM-1 expression, but treating HUVECs with cytomix caused significantly higher ICAM-1 expression even in the presence of TAK-242 (supplementary Fig. S1). When we dosed HUVECs with BEVs derived from *E. coli* strains CFT073 [WAM2267] or K1 RS218 for 16–17 h, co-treatment with TAK-242 brought ICAM-1 expression back to baseline levels (Fig. 2). Therefore, we concluded that TLR4 is the primary receptor mediating the stimulatory interactions between *E. coli*-derived BEVs and HUVECs, and it is likely that LPS is a major stimulatory component of these BEVs, consistent with findings reported in previous studies^20,41,42^. To our knowledge, this is the first demonstration that blockade of TLR4 can protect HUVECs from BEV-induced upregulation of ICAM-1.

**Fig. 2:**
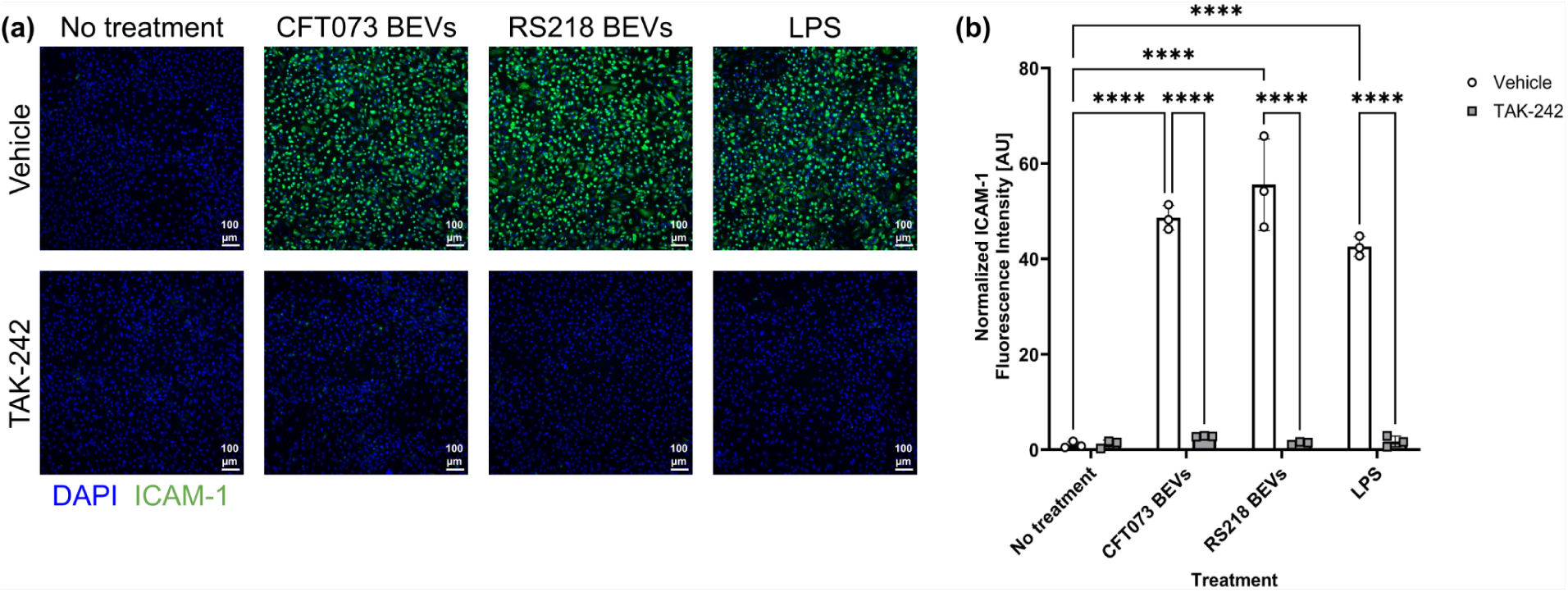
BEVs stimulate HUVECs by binding to TLR4. (**a**) Immunofluorescence images of HUVECs stained with DAPI (blue) and anti-ICAM-1 antibody (green). Cells were pre-treated with 10 µM of TLR4 inhibitor TAK-242 or a vehicle control (0.04% DMSO) for 4–5 h. They were subsequently exposed to 10 ng/ml LPS or 1E8 BEVs/ml derived from *E. coli* strains CFT073 [WAM2267] or K1 RS218 with either the TAK-242 or vehicle control for 16–17 h. (**b**) Barplot of ICAM-1 fluorescence intensity. The average fluorescence intensity of ICAM-1 in each image was background-subtracted and then divided by the number of cell nuclei in the corresponding DAPI channel. These values were then normalized to the average for the wells that received the vehicle control with no stimulatory treatment. Error bars = s.d. Performed two-way ANOVA with Tukey post-hoc test: ✱✱✱✱ = *p* < 0.0001

### Estimating minimum inhibitory concentration values

The Minimum Inhibitory Concentration (MIC) is the lowest concentration of an antimicrobial agent that inhibits visible bacterial growth. In this study, the MICs of meropenem, tobramycin, and ciprofloxacin for *E. coli* strain CFT073 [WAM2267] were determined via the broth-dilution assay, adhering to EUCAST guidelines (Table 1)^43^. The MICs for strain K1 RS218 had previously been determined by our group^18^.

**Table 1:** Comparison of MIC values between *E. coli* strains CFT073 [WAM2267] and K1 RS218 for meropenem, tobramycin and ciprofloxacin.

| *E. coli* Strain | Meropenem MIC (μg/ml) | Tobramycin MIC (μg/ml) | Ciprofloxacin MIC (μg/ml) |
| --- | --- | --- | --- |
| CFT073 [WAM2267] | 0.025 | 10 | 0.1 |
| K1 RS218 | 0.1 | 10 | 0.1 |

While RS218 and CFT073 showed similar MICs for tobramycin and ciprofloxacin, the MIC for meropenem was fourfold higher in RS218. This difference may be due to RS218’s unique virulence features, such as its K1 capsule and enhanced ability to persist within host cells, which could confer increased tolerance to cell wall–targeting antibiotics like meropenem, even in the absence of classic resistance mechanisms^25,44,45^.

### Characterization of BEVs derived from antibiotic-treated *E. coli*

*E. coli* were incubated for 3.5 h with twice the MIC (2MIC) of meropenem, tobramycin, ciprofloxacin, or no treatment (control) before BEVs were collected via differential centrifugation. After resuspending the BEVs in PBS, they were quantified by nanoparticle tracking analysis (NTA). Three biological replicates were collected for each examined condition. We observed that the average concentration of recovered BEVs from the control conditions was 3.91E10 ± 1.81E9 particles/ml for strain CFT073 and 1.71E11 ± 8.69E9 particles/ml for strain RS218 (Fig. 3a). In other words, there was nearly an order of magnitude more BEVs recovered from the RS218 cultures compared to the other strain. This trend remained true for all of the antibiotic-treated conditions: meropenem treatment resulted in recovery of 1.48E10 ± 3.29E8 CFT073 BEVs/ml and 2.52E11 ± 1.05E10 RS218 BEVs/ml, tobramycin treatment resulted in recovery of 4.53E10 ± 1.58E9 CFT073 BEVs/ml and 1.98E11 ± 1.02E10 RS218 BEVs/ml, and ciprofloxacin treatment resulted in recovery of 2.73E10 ± 1.08E9 CFT073 BEVs/ml and 1.89E11 ± 1.06E10 RS218 BEVs/ml (Fig. 3a). Notably, meropenem treatment resulted in a significant increase in production of BEVs by strain RS218 over the control condition, suggesting a potential link between cell wall-targeting antibiotics and increased vesiculation. However, there was no significant change in BEV production by strain CFT073 when exposed to meropenem and, in fact, there was a non-significant trend towards decreased BEV production under these conditions. This may highlight another strain-specific difference in behavior, or could be related to the different meropenem MICs that were identified for the two strains.

**Fig. 3:**
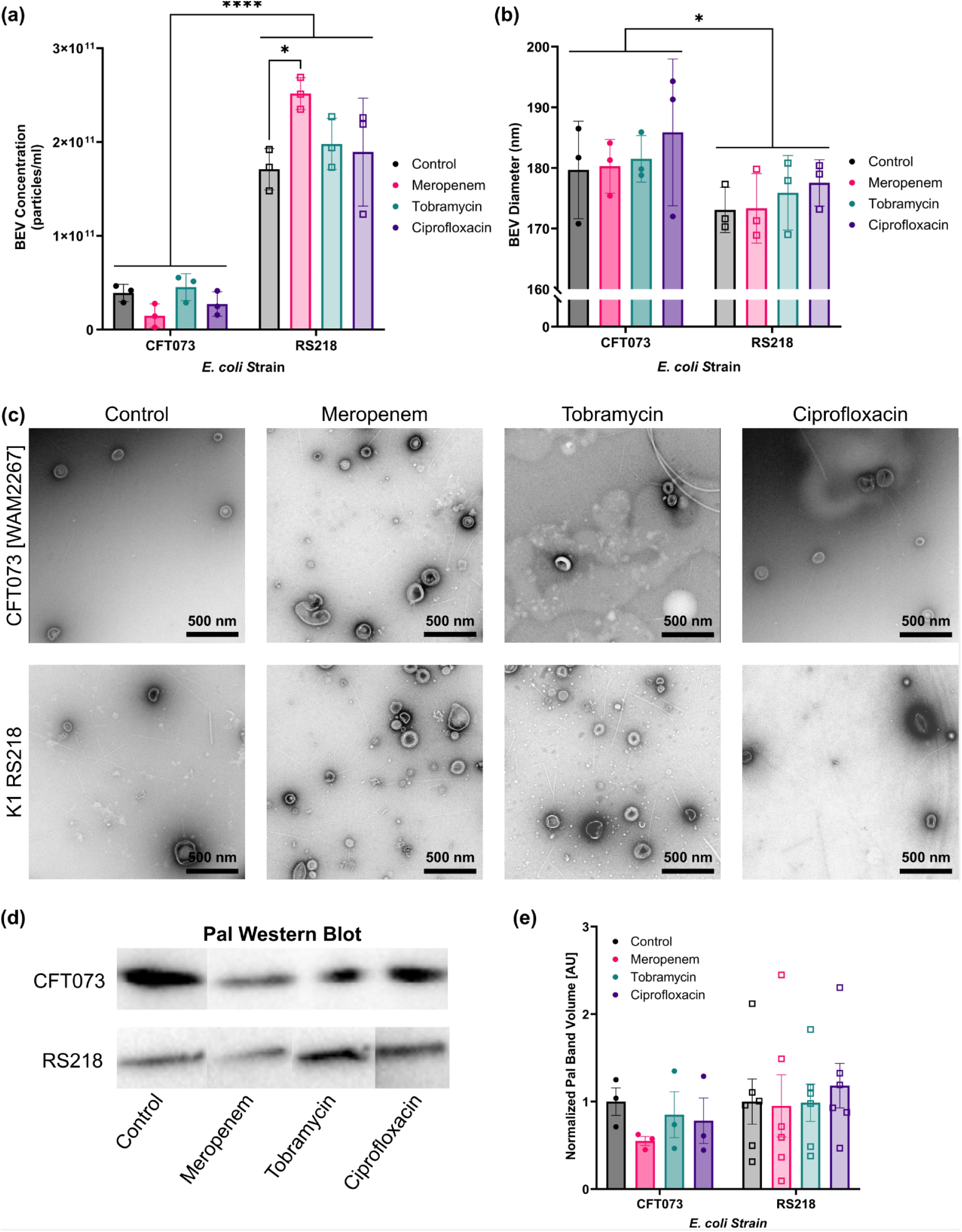
*E. coli* demonstrate strain-specific changes in BEV production after antibiotic treatment. (**a**) Barplot of BEV concentrations as measured by nanoparticle tracking analysis (NTA). Performed two-way ANOVA with Tukey post-hoc test: ✱ = p < 0.05, ✱✱✱✱ = p < 0.0001. Error bars = s.d. (**b**) Barplot of BEV diameters as measured by NTA. Performed two-way ANOVA with Tukey post-hoc test and found a significant effect due to the different *E. coli* strains: ✱ = p < 0.05. Error bars = s.d. (**c**) Transmission electron microscopy images of BEVs derived from *E. coli* strains CFT073 [WAM2267] and K1 RS218 treated with various antibiotics or no antibiotic (control). (**d**) Western Blot of Pal in equal volumes of the BEV samples. All of the CFT073 blots have been rearranged from a single gel image, and all of the RS218 blots have been rearranged from a single gel image. For the original full gel images, see supplementary Fig. S2. (**e**) Barplot of Pal band densities from Western Blot images. Error bars = s.e.m.

BEV size measurements were also obtained from NTA (Fig. 3b). There were no significant changes in particle size due to antibiotic treatment. However, there was a significant difference in the sizes of BEVs produced by the two *E. coli* strains; the average diameter of BEVs derived from CFT073 was 181.8 ± 2.063 nm while the average for RS218 was 175.0 ± 1.346 nm. This size discrepancy was not necessarily apparent in all of the transmission electron microscopy (TEM) images that were captured for the various BEV preparations (Fig. 3c). The TEM images did, however, agree with the NTA data concerning the increase in vesicle production when strain RS218 was treated with meropenem. These TEM images also confirmed that the recovered particles featured the typical morphology of BEVs^46^.

Western Blot of equal volumes of the BEV preparations confirmed the presence of peptidoglycan-associated lipoprotein (Pal) in all samples (Fig. 3d). For strain CFT073, there was a non-significant decrease in Pal content due to meropenem treatment (Fig. 3e). This corresponded with the non-significant drop in the concentration of BEVs when the CFT073 *E. coli* were treated with meropenem as measured by NTA. However, the meropenem-induced increase in production of BEVs by strain RS218 did not cause a noticeable change in Pal band density in the Western Blots. It is possible that less Pal was packaged in RS218-derived BEVs due to meropenem treatment, which could be another strain-specific behavioral difference.

### Some, but not all, antibiotic treatments at 2MIC reduced the bulk stimulatory capacity of *E. coli*-derived BEVs

To explore the influence of antibiotic-induced BEVs on endothelial activation, we measured ICAM-1 expression in HUVECs treated with BEVs derived from *E. coli* strains CFT073 [WAM2267] and K1 RS218 that had been exposed to different antibiotics. HUVECs were incubated with BEVs for 16–17 hours, after which ICAM-1 expression was assessed using immunofluorescence staining (Fig. 4a,c). For each strain, BEVs derived from the control condition (no antibiotic treatment) were diluted to 1E8 particles/ml. The same dilution factor was then applied to all of the BEVs obtained from the antibiotic-treated conditions such that the HUVECs were treated with equal volumes of each BEV preparation. Quantitative analysis of the fluorescence intensity revealed that BEVs from *E. coli* strain CFT073 [WAM2267] that had been treated with meropenem or tobramycin resulted in significantly lower ICAM-1 expression compared to the control, indicating a lower stimulatory potential (Fig. 4b). CFT073 BEVs from the ciprofloxacin conditions also demonstrated a non-significant trend towards decreased stimulatory capacity. BEVs from strain K1 RS218 that had been treated with tobramycin exhibited significantly lower ICAM-1 expression than the control, meropenem, and ciprofloxacin conditions (Fig. 4d). However, there were no significant differences between these other conditions. These findings suggest that meropenem exposure alters the stimulatory capacity of BEVs in a strain-dependent manner, potentially due to changes in BEV composition or because of the differences in BEV production demonstrated by the two strains during meropenem treatment (Fig. 3a). Conversely, tobramycin seems to consistently reduce the stimulatory potential of BEVs regardless of *E. coli* strain. The reduced ability of BEVs from antibiotic-treated bacteria to upregulate ICAM-1 may have implications for modulating endothelial inflammation during infections and antibiotic therapy. Furthermore, it is notable that ciprofloxacin treatment did not significantly lower the stimulatory capacity of BEVs from either strain. This indicates that, while ciprofloxacin may be effective at treating bacterial infections, the BEVs produced by the inhibited or dying bacteria may still demonstrate equal stimulatory effects to BEVs secreted by an untreated *E. coli* infection.

**Fig. 4:**
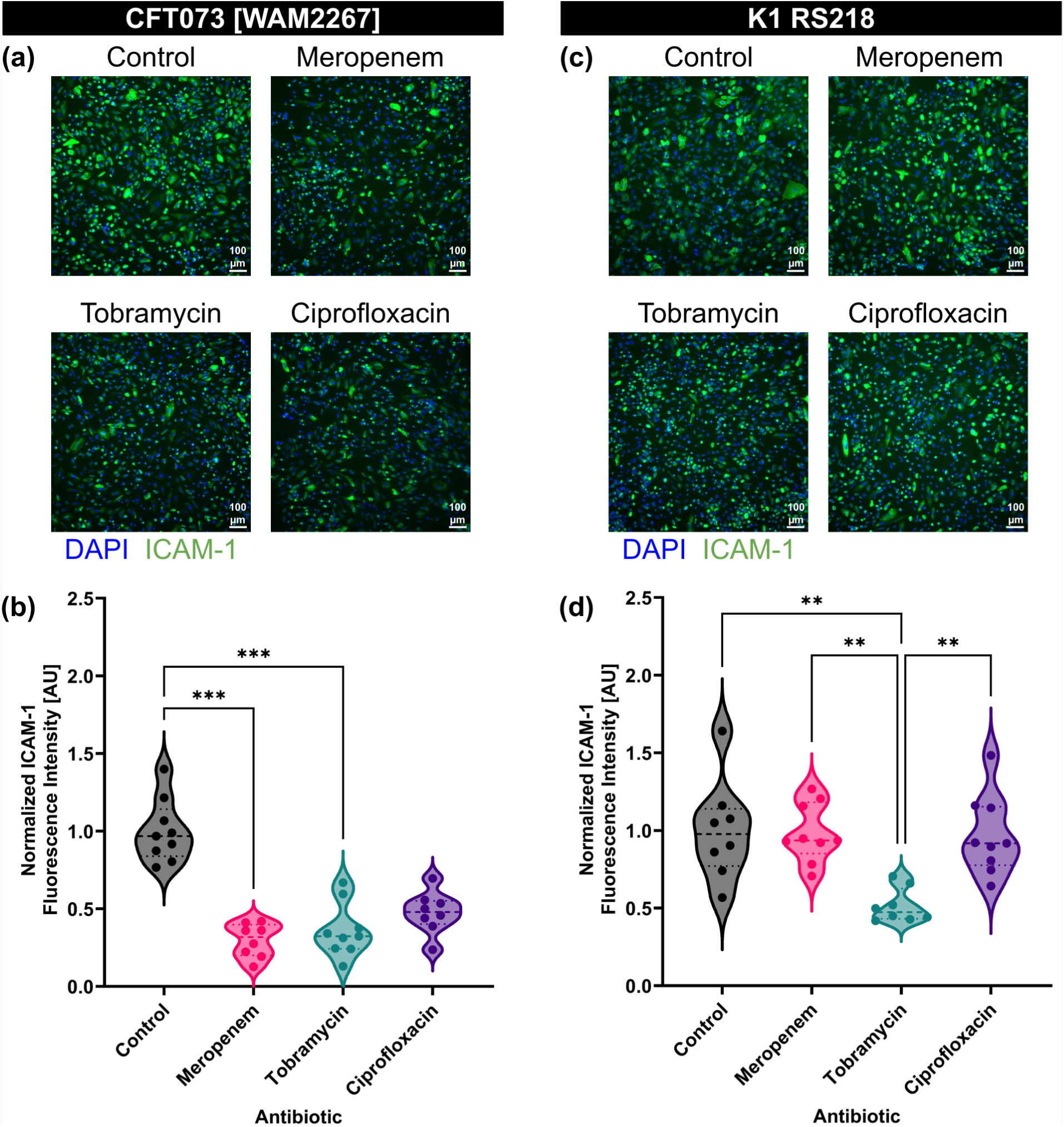
Treatment of *E. coli* with antibiotics can lower the stimulatory capacity of equal volume BEV preparations. (**a**) Immunofluorescence images of HUVECs stained with DAPI (blue) and anti-ICAM-1 antibody (green). Cells were treated with BEVs derived from *E. coli* CFT073 [WAM2267] that had been treated without antibiotics (control) or with 2MIC of meropenem, tobramycin, or ciprofloxacin for 16–17 h. The control BEVs were diluted to 1E8 particles/ml, and the same dilution factor was applied to the other experimental conditions to ensure that cells were treated with equal volumes of BEVs. (**b**) Violin plot of ICAM-1 fluorescence intensity from HUVECs treated with BEVs derived from *E. coli* strain CFT073 [WAM2276]. The average fluorescence intensity of ICAM-1 in each image was divided by the number of cell nuclei in the corresponding DAPI channel. The baseline ICAM-1 fluorescence intensity of untreated HUVECs was then subtracted, and the resulting values were normalized to the mean of the control condition. Performed Kruskal-Wallis with Dunn’s correction: ✱✱✱ = *p* < 0.001 (**c**) Immunofluorescence images of HUVECs stained with DAPI (blue) and anti-ICAM-1 antibody (green). Cells were treated with BEVs derived from *E. coli* K1 RS218 that had been treated without antibiotics (control) or with 2MIC of meropenem, tobramycin, or ciprofloxacin for 16–17 h. The control BEVs were diluted to 1E8 particles/ml, and the same dilution factor was applied to the other experimental conditions to ensure that cells were treated with equal volumes of BEVs. (**d**) Violin plot of ICAM-1 fluorescence intensity from HUVECs treated with BEVs derived from *E. coli* strain K1 RS218. The average fluorescence intensity of ICAM-1 in each image was divided by the number of cell nuclei in the corresponding DAPI channel. The baseline ICAM-1 fluorescence intensity of untreated HUVECs was then subtracted, and the resulting values were normalized to the mean of the control condition. Performed Kruskal-Wallis with Dunn’s correction: ✱✱✱ = *p* < 0.001

### Treating CFT073 *E. coli* with meropenem increases the stimulatory capacity of individual BEVs

To investigate the impact of antibiotic treatment on the stimulatory potential of individual BEVs, we standardized the BEV concentrations to 1E8 BEVs/ml for all conditions before exposing them to HUVECs. Immunofluorescence staining for ICAM-1 revealed strain-dependent differences in endothelial activation (Fig. 5a,c).

**Fig. 5:**
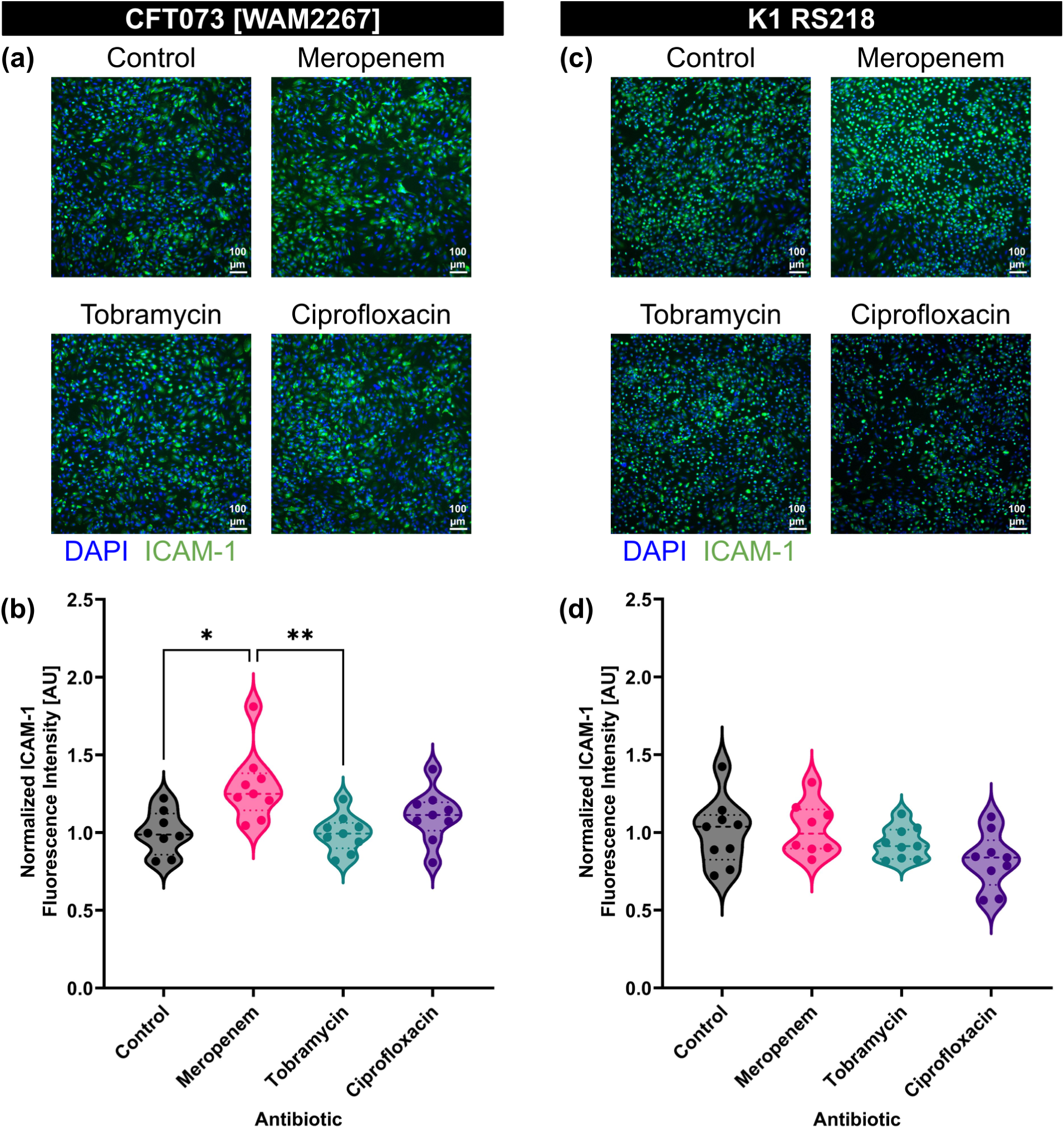
Meropenem treatment increases the potency of individual BEVs derived from *E. coli* strain CFT073 [WAM2267]. (**a**) Immunofluorescence images of HUVECs stained with DAPI (blue) and anti-ICAM-1 antibody (green). Cells were treated with BEVs derived from *E. coli* CFT073 [WAM2267] that had been treated without antibiotics (control) or with 2MIC of meropenem, tobramycin, or ciprofloxacin for 16–17 h. All treatment concentrations were standardized to 1E8 BEVs/ml. (**b**) Violin plot of ICAM-1 fluorescence intensity from HUVECs treated with BEVs derived from *E. coli* strain CFT073 [WAM2276]. The average fluorescence intensity of ICAM-1 in each image was divided by the number of cell nuclei in the corresponding DAPI channel. The baseline ICAM-1 fluorescence intensity of untreated HUVECs was then subtracted, and the resulting values were normalized to the mean of the control condition. Performed Kruskal-Wallis with Dunn’s correction: ✱ = *p* < 0.05, ✱✱ = *p* < 0.01. (**c**) Immunofluorescence images of HUVECs stained with DAPI (blue) and anti-ICAM-1 antibody (green). Cells were treated with BEVs derived from *E. coli* K1 RS218 that had been treated without antibiotics (control) or with 2MIC of meropenem, tobramycin, or ciprofloxacin for 16–17 h. All treatment concentrations were standardized to 1E8 BEVs/ml. (**d**) Violin plot of ICAM-1 fluorescence intensity from HUVECs treated with BEVs derived from *E. coli* strain K1 RS218. The average fluorescence intensity of ICAM-1 in each image was divided by the number of cell nuclei in the corresponding DAPI channel. The baseline ICAM-1 fluorescence intensity of untreated HUVECs was then subtracted, and the resulting values were normalized to the mean of the control condition. Performed Kruskal-Wallis with Dunn’s correction and found no significant differences.

For CFT073 [WAM2267]-derived BEVs, ICAM-1 expression was higher when BEVs were obtained from bacteria treated with meropenem, while BEVs from tobramycin- and ciprofloxacin-treated bacteria induced ICAM-1 expression at levels similar to the control (Fig. 5b). This suggests that meropenem exposure may enhance the stimulatory potential of BEVs, whereas tobramycin and ciprofloxacin do not substantially alter the BEVs’ effect on endothelial activation.

In contrast, for K1 RS218-derived BEVs, ICAM-1 expression levels remained comparable across treatment groups, indicating that the stimulatory potential of BEVs from this strain was not substantially affected by antibiotic exposure (Fig. 5d). These findings highlight potential strain-specific differences in how antibiotics influence BEV cargo and their downstream effects on endothelial cells.

### Protein profiling

To investigate why meropenem treatment impacted the stimulatory capacity of BEVs derived from *E. coli* strain CFT073 [WAM2267], we performed mass spectrometry and looked for changes in BEV cargo composition. BEVs isolated from *E. coli* treated with meropenem, tobramycin, ciprofloxacin, or the untreated control were all included for comparison (Fig. 6). This analysis allowed us to identify changes in the protein cargo of BEVs in response to different antibiotic classes, revealing potential alterations in bacterial stress responses, virulence factor secretion, and membrane-associated proteins. By comparing the proteomic profiles across conditions, we aimed to determine how antibiotic-induced changes in BEV content could influence host-pathogen interactions, particularly endothelial activation and inflammation.

**Fig. 6:**
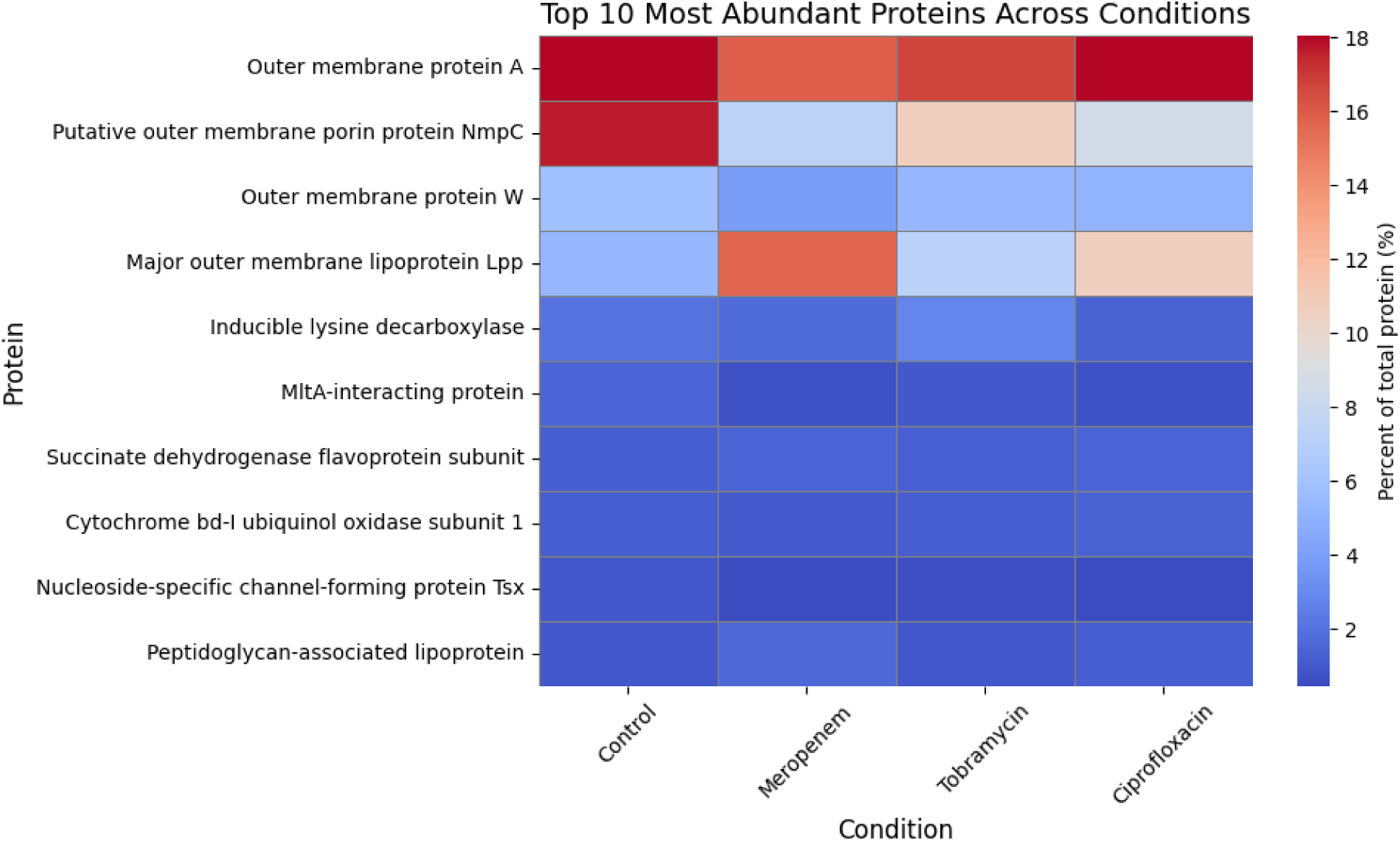
Mass spectrometry-based heatmap of the top 10 most abundant BEV proteins across conditions showcases differences in NmpC and Lpp levels. Relative protein abundance (% of total protein) in BEVs from *E. coli* exposed to meropenem, tobramycin, ciprofloxacin, or no antibiotic (control). Data reflect the top 10 most abundant proteins identified by mass spectrometry. Outer membrane proteins dominated across all conditions, with antibiotic-specific shifts observed.

Among the most abundant proteins detected across all conditions were outer membrane proteins (OMPs), including OmpA, NmpC, and major outer membrane lipoprotein Lpp, which are known to play critical roles in vesicle structure, host interaction, and immune activation^47^. OmpA remained consistently enriched across all conditions, indicating its stable incorporation into BEVs regardless of antibiotic treatment and underscoring its structural and immunogenic significance^48^. In contrast, NmpC, a porin involved in nutrient uptake and stress adaptation^49^, was highly enriched in control conditions but markedly reduced following antibiotic treatment, possibly indicating downregulation of porin expression to limit antibiotic infiltration into the *E. coli*^50^. Interestingly, lipoprotein Lpp, which tethers the outer membrane to the peptidoglycan layer, was notably increased in BEVs following meropenem treatment and to a lesser degree by ciprofloxacin treatment^51,52^. These findings indicate antibiotic-dependent modulation of the proteins present in *E. coli*-derived BEVs, and it is possible that the altered levels of Lpp could explain the stronger stimulatory potential of individual BEVs derived from meropenem-treated *E. coli*.

## Discussion

In this work, we examined how treatment of two strains of *E. coli* with various antibiotics affected the production and stimulatory characteristics of BEVs. The results showcased differences between two strains of the same species, which may help inform clinical treatments to mitigate sepsis stemming from either UTIs or meningitis. By modeling the interactions between BEVs and human endothelial cells, we recreated a potentially important component of sepsis that can promote endothelial inflammatory activation and thereby attract immune cells to the vascular wall. While bacteria are often responsible for initiating sepsis, the negative outcomes associated with this disease state are more commonly attributed to the host immune response including increased production of pro-inflammatory cytokines^2^. However, the interaction between bacterial products and the endothelium should not be ignored as it may play a role in destabilizing vital regions of the vasculature such as the blood-brain barrier, leading to neurological changes that could continue to impact sepsis survivors, even following clearance of the bacterial infection^6,53^.

We found that HUVECs exhibited a dose-dependent increase in ICAM-1 expression following exposure to *E. coli*-derived BEVs. This underscores the potential risks of treating bacteria with antibiotics that could promote increased BEV production. However, we also found that the increase in BEV release by meropenem-treated *E. coli* from strain K1 RS218, which likely occurred due to meropenem’s disruption of tethers between the outer membrane and the bacterial cell wall, did not exhibit increased stimulatory effects on the HUVECs^17^. In this instance, the relationship between higher doses of BEVs and ICAM-1 expression was not one-to-one, indicating that there was either a decrease in the stimulatory capacity of the BEVs from the meropenem condition or that differences in BEV potency may only be apparent when concentrations differ by at least an order of magnitude. Similarly, ICAM-1 expression appeared to be inversely related to the dose of BEVs derived from tobramycin-treated *E. coli*. This directly contradicts the dose-response data, but a potential explanation stems from the fact that tobramycin itself has been demonstrated to have anti-inflammatory effects^54^. While we suspect that the majority of free tobramycin was removed from the BEV preparations following ultracentrifugation, there is a possibility that a small amount of tobramycin remained behind or that it had been encapsulated within the BEVs. This could account for the paradoxically lower ICAM-1 response when the HUVECs were treated by a larger number of BEVs.

One other important factor that significantly influenced the potency of the BEVs was the source strain. Despite the fact that CFT073 [WAM2267] and K1 RS218 are the same bacterial species, BEVs from the former strain were capable of increasing HUVEC ICAM-1 expression at a ten-fold lower concentration than BEVs from the latter strain. This could reflect the ability of K1 RS218 to evade the host inflammatory response, for example by producing the K1 capsule, which resembles structures present on neurons and immune cells and helps to disguise the invading pathogen^24^. It is conceivable that BEVs produced by this strain would also possess surface motifs that could act as camouflage when the particles remain below a certain concentration threshold. There are also a number of gram-positive and gram-negative bacteria that express anti-phagocytic proteins, and packaging of anti-autophagic cargo into BEVs of *E. coli* strain SP15 has also been observed^15,16^. Such species- and strain-specific differences in defenses and host evasion tactics could certainly account for the different levels of ICAM-1 upregulation observed in the present study. With that said, it is curious that K1 RS218 produced an order of magnitude more BEVs than CFT073 [WAM2267]. Genetic differences can result in altered BEV production by *E. coli*, with *nlpI* and *degP* mutants demonstrating hypervesiculation and *nlpA* mutants demonstrating hypovesiculation^55,56^. It is likely that the pronounced disparity in BEV production observed in this study is due to the strain-specific genomes, though additional work may be required to determine which particular mutations are directly responsible. Additionally, differences in BEV production might occur if the *E. coli* were co-cultured directly with the human cells. It is clear that environmental stressors such as iron deficiency, which occurs in the host environment, can impact vesiculation from bacteria^57^.

Another strain-specific difference was the size of the secreted BEVs as detected by NTA. CFT073 [WAM2267] produced BEVs with an average diameter greater than the K1 RS218-derived BEVs. This could also result in a larger interaction area between individual vesicles and the endothelium, potentially leading to increased presentation of surface LPS and other inflammatory factors to the endothelial cells. This mechanism could also explain the measured differences in BEV potency. Some of the trends from the NTA data were also apparent in the TEM images. A subset of these images included flagella that were co-isolated with BEVs, which was a limitation of the ultracentrifugation technique used in this study. It is worth noting that the presence of flagella could also have an impact on the values recorded from NTA if any were mistakenly counted as individual particles. Flagella can also impact inflammation by stimulating cells through TLR5, but based on our TLR inhibition studies, it seemed that this pathway was not strongly engaged in our HUVECs^58^. Therefore, although our ultracentrifugation-based BEV isolation was not completely pure, the contribution of flagella was negligible in the studies with endothelial cells and the observed results can be attributed to the BEVs.

TLRs are type 1 transmembrane proteins that play a key role in the immune system by recognizing various molecular signals, including pathogen-associated signals, damage-associated molecular patterns, such as calprotectin, and pathogen-associated molecular patterns, such as the aforementioned flagellin and LPS^59^. These receptors are found on immune cells, such as dendritic cells and macrophages, and non-immune cells, such as fibroblasts and endothelial cells. Upon ligand binding, TLRs initiate a signaling cascade primarily through adaptor molecules such as MyD88 or TRIF, leading to the activation of transcription factors like NF-κB^60^. This results in the upregulation of pro-inflammatory cytokines and type I interferons, which coordinate an immune response to combat infection or cellular damage^27^. TLR4 recognizes LPS from gram-negative bacteria, leading to the activation of both anti-inflammatory and pro-inflammatory pathways^61^. The engagement of TLR4 by BEVs results in the upregulation of inflammatory markers such as ICAM-1, CXCL10, and IL-6, which are crucial in mediating endothelial cell activation and the subsequent inflammatory response^20,41^.

TAK-242 is a specific inhibitor of TLR4 and has previously been demonstrated to have no functional effect on other TLRs^62^. It functions by binding to TLR4’s intracellular domain to disrupt the interactions with the MyD88 and TRIF adaptor molecules and thereby prevent the pro-inflammatory signaling cascade^62,63^. In the present study, we validated TAK-242’s specificity for inhibiting TLR4 by examining its ability to prevent endothelial ICAM-1 upregulation during LPS treatment but not during treatment with a pro-inflammatory cytomix consisting of equal parts TNF-α, IL-1β, and IFN-γ. These three cytokines engage receptors other than TLR4 and thus upregulate endothelial ICAM-1 expression via alternate pathways. Interestingly, there was a significant decrease in ICAM-1 expression when TAK-242 was administered with the cytomix, though the ICAM-1 signal remained far above basal levels. Even though the cytomix components promote endothelial activation via non-TLR4 receptors, it is possible for all three of these cytokines together to induce upregulation of TLR4^64–66^. TLR4 overexpression, in turn, can cause an increase in ICAM-1 expression by endothelial cells^67^. It is therefore conceivable that the change in ICAM-1 signal during TAK-242 cotreatment with cytomix was due to inhibition of excess TLR4, but the fact that ICAM-1 expression remained elevated indicated that the other pro-inflammatory pathways were still engaged.

In our experiments with BEVs, TAK-242 treatment prevented increases in ICAM-1. This indicated that the primary component of *E. coli* BEVs responsible for stimulating endothelial cells was likely LPS since the inhibited receptor, TLR4, is responsible for sensing LPS. Laakmann et al. similarly demonstrated that lung endothelial cells could be protected from the pro-inflammatory effects of BEVs derived from gram negative bacteria by neutralizing LPS with polymyxin B^20^. Ho et al. showed that BEVs derived from *Porphyromonas gingivalis*, another gram-negative species, could promote TLR4 upregulation in HUVECs while Li et al. demonstrated that the same was true for *E. coli*-derived BEVs^68^. However, the results in the present study are, to our knowledge, the first to demonstrate that TLR4 blockade could protect HUVECs from the stimulatory effects of BEVs. Additional findings across the literature have made inhibition of LPS-TLR4 signaling an attractive goal for sepsis mitigation. In animal models, treatment with TAK-242 prevented LPS-induced damage to kidneys, lungs, and skeletal muscle^63,69,70^. Unfortunately, TAK-242 was not an effective sepsis treatment in human phase III clinical trials^28^. This may be due to the inherent complexity of sepsis, though it is worth noting that these trials did not include TAK-242 pre-treatment since the patients were already suffering from sepsis prior to TAK-242 administration. Additionally, the main readouts from these trials were measurements of serum IL-6 levels and mortality rates rather than endothelial ICAM-1 upregulation, so it is possible that TLR4 inhibition could still prove useful in combination with other therapies^28^. Significantly, BEVs possess other cargo that is known to have pro-inflammatory properties, including outer membrane protein A and peptidoglycan-associated lipoprotein^17,71^. Even if these factors did not contribute significantly to ICAM-1 expression on endothelial cells in the present study, it is possible that they would have a substantial impact on immune cells and other cell types even if TLR4 were inhibited.

One other example of BEV cargo that could serve as a stimulant is major outer membrane lipoprotein Lpp, also referred to as Braun’s lipoprotein. Mass spectrometry revealed a large increase in the proportion of Lpp in BEVs derived from CFT073 [WAM2267] *E. coli* due to treatment with meropenem. This makes sense since β-lactam antibiotics such as meropenem act by disrupting the bacterial cell wall, which could untether the Lpp connecting the peptidoglycan layer with the outer membrane^35,72^. The altered abundance of this protein suggests antibiotic-specific modulation of BEV cargo, likely linked to membrane stress responses and envelope integrity maintenance mechanisms^18^. The greater Lpp content in BEVs derived from meropenem-treated CFT073 *E. coli* could explain the significant increase in ICAM-1 expression that occurred when HUVECs were exposed to these BEVs since Lpp has previously been observed to promote endothelial pro-inflammatory activation^73^. Lpp has also been shown to cause inflammation *in vitro* and *in vivo* via interactions with TLR2^74^. A future extension to the present study would be to test TLR2 blockade to see if this eliminates the enhanced stimulatory effects of the BEVs derived from meropenem-treated CFT073 *E. coli*. Although these meropenem-associated BEVs were more stimulatory than the other CFT073 BEV preparations, this effect was only apparent when HUVECs were treated with equal concentrations of the BEV preparations. Overall, meropenem treatment significantly reduced the ability of secreted BEVs to cause ICAM-1 upregulation, probably because of a non-significant reduction in the production of BEVs by CFT073 *E. coli*. This once again emphasizes the importance of the size of the BEV dose experienced by the endothelium.

Another interesting change in BEV protein cargo that was observed in our mass spectrometry results was the reduction of putative outer membrane porin protein NmpC in all antibiotic-treated conditions compared to the untreated control. Porins serve as channels for passive diffusion through the outer membrane and thus could act as conduits for antibiotics to access the interior of the bacteria. Mutations in porin-encoding genes as well as changes in porin expression have been observed as a means for bacteria to resist antibiotic treatment^50^. Since many of our BEVs may have originated from the outer membrane, the reduced abundance of NmpC in these vesicles probably stems from changes in the parent bacteria outer membrane that were made to prevent antibiotics from entering the *E. coli*.

This work was largely focused on endothelial expression of ICAM-1, which primes the endothelial cells for interactions with passing immune cells. In the future, it may be interesting to see whether BEVs derived from bacteria treated with varying antibiotics may also have differing effects on these downstream endothelial-immune interactions. In addition to dosing the endothelium, the effect of antibiotic-induced BEVs on immune cell morphology, migratory response, cytokine production, and expression of inflammatory markers could help inform researchers and clinicians about how antibiotic treatment modulates key processes during sepsis. A limitation of this work is that we only included one representative from each of three antibiotic classes: β-lactams, aminoglycosides, and fluoroquinolones. In order to generalize these results, similar experiments should be performed with other examples of these antibiotic classes. Furthermore, only a single bacterial species was examined. It is possible that strain-specific differences, such as the ones determined in this study, may occur across many sepsis-associated bacterial species. Much additional work can be performed in order to create a more complete understanding of the relationships between antibiotics, bacterial strains, and BEV production and pro-inflammatory capacity.

In summary, our findings highlight that antibiotic treatment not only influences the quantity of BEVs released by different *E. coli* strains but also alters their capacity to stimulate endothelial cells. These antibiotic-driven changes in BEV production and composition significantly impact endothelial activation, including ICAM-1 upregulation, suggesting a previously underappreciated link between antimicrobial therapy and host vascular responses.

## Materials and methods

### Cell culture

Pooled primary HUVECs were obtained from ATCC (PCS-100-013), Lonza (C2519A), and Thermo Fisher (C01510C). Cells were maintained in Human Large Vessel Endothelial Cell Medium (Gibco, M200500) supplemented with 2% Large Vessel Endothelial Supplement (Gibco, A1460801) and 1% Penicillin/Streptomycin, which was exchanged every 2–3 days. HUVECs were incubated at 37 °C, 5% CO_2_. For experiments, cells were seeded on 96-well plates at a density of approximately 40,000 cells/cm^2^ and were used up to passage 9.

### BEV isolation

*E. coli* were cultured on Luria-Bertani (LB) agar at 37 °C overnight; colonies from the plate were used to inoculate a 50 ml growth in LB broth, which was cultured overnight at 37 °C, shaking at 160 rpm. This small culture was divided in half to inoculate two larger cultures of 200 ml each, which were grown to log phase (optical density at 600 nm ∼0.8). These cultures were split into 49.5 ml aliquots in sterilized 125 ml flasks. All antibiotic solutions were prepared at 2MIC and then added into their corresponding flasks (500 µl of antibiotic solution + 49.5 mL of media). The *E. coli* cultures were incubated with antibiotics (or no antibiotic, for control) for 3.5 h (37 °C, 180 rpm). Afterward the samples were first centrifuged at 5,000×g for 15 min to pellet the *E. coli*. Then the supernatant was passed through a 0.45 µm syringe filter and ultracentrifuged at 100,000×g for 2 h to pellet the BEVs. Pellets were resuspended in 400 µL of PBS so that the final BEV preparations were brought to 125 times their original concentration, and were either kept at 4 °C for less than 1 month or at −20 °C until needed for experiments. To confirm that no live *E. coli* were co-isolated with the BEVs, samples were mixed in cell culture media and inspected for signs of growth after overnight incubation at 37 °C, 5% CO_2_. As a control, sterile LB broth alone was subjected to nanoparticle tracking analysis (see below), revealing that negligible vesicle-sized particles were present in the absence of bacteria.

### Endothelial stimulation and immunofluorescent labeling

BEVs or purified LPS (Sigma-Aldrich, L3012-5MG) were diluted in cell culture media and then added to confluent HUVECs in a 96-well plate. Treatment lasted 16–17 h at 37 °C, 5% CO_2_. Cells were treated with anti-ICAM-1 antibody (Biolegend, 353102) diluted 1:100 in cell culture media for 15 min, then washed with PBS, and fixed with 4% formaldehyde for 15 min. The fixed cells were subsequently washed three times with PBS, blocked with 10% normal goat serum (Thermo Fisher, 50062Z) for 30 min at room temperature, then stained with a secondary goat anti-mouse IgG antibody conjugated to Alexa Fluor 488 (Thermo Fisher, A-11001) diluted 1:200 in 10% normal goat serum for 2 h in the dark at room temperature. Following this incubation, cells were washed with PBS and their nuclei were labeled with DAPI (Thermo Fisher, D1306) diluted 1:400 in deionized water for 3 min at room temperature prior to three final washes with PBS. Samples were imaged on a Leica DMI6000 B fluorescence microscope. To quantify ICAM-1 labeling, background signal was subtracted from the average fluorescence measured from GFP channel images. This background-subtracted average was then divided by the number of nuclei counted in DAPI channel images to yield an estimate of the amount of ICAM-1 fluorescence signal per individual cell. To determine the BEV concentration in terms of BEVs/HUVEC, the total number of BEVs per well was calculated based on the volume added to each well of the 96-well plate (100 µl). This was divided by the estimated number of HUVECs in each well, which was found by taking the number of nuclei counted in DAPI channel images, calculating the average number of cells per unit area, and then multiplying this by the known area of the well.

### Blockade of TLR4

To investigate the role of TLR4 in BEV-induced inflammation, we used a specific antagonist, TAK-242, to block the TLR4 receptor. HUVECs were pre-treated with 10 µM TAK-242 or a 0.04% DMSO vehicle control for 4–5 h prior to introducing 1E8 BEVs/ml or 10 ng/ml LPS with additional TAK-242 or vehicle control for 16–17 h. The cells were subsequently stained for ICAM-1 (see above). To confirm that TAK-242 did not inhibit the ability of other receptors to cause increased ICAM-1 expression, HUVECs were treated with 500 pg/ml pro-inflammatory Cytomix consisting of equal parts TNF-α, IL-1β, and IFN-γ.

### Determination of minimum inhibitory concentration (MIC)

The minimum inhibitory concentration (MIC) of each antibiotic was determined using the broth-dilution method according to EUCAST guidelines, with a modification to use LB media to match the conditions of all subsequent *E. coli* culture experiments^75^. *E. coli* cultures were grown overnight in LB broth and inoculated into test tubes containing fresh LB broth with serially diluted antibiotics. Eight serial 1:2 dilutions were prepared for each antibiotic to create a concentration range. An antibiotic-free control was included to ensure optimal bacterial growth conditions. The cultures were incubated overnight at 37 °C with shaking at 160 rpm, and bacterial growth was assessed the next day by examining the tubes for turbidity, which indicated visible growth. The MIC was defined as the lowest antibiotic concentration that inhibited visible growth, keeping the broth clear. For subsequent experiments, *E. coli* cultures were treated with antibiotics standardized at 2MIC to enable comparative analysis across different antibiotics.

### Nanoparticle tracking analysis (NTA)

Nanoparticle size distributions and concentrations were assessed using a NanoSight NS300 (Malvern Panalytical, Malvern, United Kingdom) equipped with a sCMOS camera, 532 nm green laser, and a 565 nm long pass filter. Multiple sample dilutions were created in PBS to ensure that at least one dilution would fall into the accurate concentration detection range between 1E8–1E9 particles/ml. 3 videos of each sample were collected for 30 seconds each, with a camera level of 15 and detection threshold of 5. Diffusion coefficients were calculated based on the particle tracks, which allowed determination of the particle size distribution. Average particle concentrations were reported after taking the sample dilution factors into account.

### Transmission electron microscopy (TEM)

The BEVs were isolated from cultures grown with 2MIC of meropenem, tobramycin, ciprofloxacin, and no antibiotic (control). 200 mesh copper grids coated with formvar/carbon film were glow discharged for 30s at 30 mA in a PELCO Easiglow prior to 3 μL of the liquid BEV sample being applied for 30s. Excess sample was wicked away and grids were exposed to three 15 μL washes with molecular grade water prior to negative staining with two applications of 10 μL filtered 0.75% uranyl formate, with wicking of excess fluid using hardened Whatman 50 filter paper, between steps. The grids were allowed to dry prior to examination on a Talos 120C transmission electron microscope equipped with a CETA 16 megapixel camera (Thermo Fisher) for image capture using TIA (Thermo Fisher). For all of the samples, 7-8 images at 28k magnification were taken in at least three distinct grid squares.

### Western blot

Samples were analyzed using standard SDS-PAGE (4-16% bis acrylamide, Precast gels, VWR) and immunoblotting techniques. Equal volumes of all samples were loaded in each well. For immunoblotting, we used anti-Pal antisera from mice inoculated with purified recombinant non-lipidated Pal protein (∼21 kDa; contains a 6xHis-tag for purification; Rochester General Hospital). Immunoblots were developed using SuperSignalTM West Femto Maximum Sensitivity Substrate (ThermoFisher) and imaged using the BioRad ChemiDoc Imaging System. Band volumes were quantified using Biorad’s Quantity One software package and normalized to the control condition for each respective bacteria strain.

### Mass spectrometry

Protein samples were loaded onto a 4–12% SDS-PAGE gel and resolved for approximately 5 minutes until a ∼10 mm band formed. After staining with SimplyBlue SafeStain (Invitrogen) and washing overnight in water, the gel band was excised, diced into 1 mm cubes, de-stained, reduced with dithiothreitol (Sigma), alkylated with iodoacetamide (Sigma), and dehydrated with acetonitrile. Trypsin (Promega) was reconstituted to 10 ng/µL in 50 mM ammonium bicarbonate and added to the gel pieces. After 30 min at room temperature, additional ammonium bicarbonate was added to fully submerge the gel, and the samples were incubated at 37 °C overnight. Peptides were extracted the following day using 50% acetonitrile and 0.1% trifluoroacetic acid (TFA), dried using a CentriVap concentrator (Labconco), desalted with homemade C18 spin columns, dried again, and reconstituted in 0.1% TFA. Peptide concentrations were measured using a fluorometric peptide assay (Thermo Fisher). Peptides were then separated by liquid chromatography using a Vanquish Neo UHPLC system (Thermo Fisher) and analyzed on an Orbitrap Astral mass spectrometer (Thermo Fisher) equipped with an Easy-Spray ion source operating at 2 kV. Samples were first trapped on a 75 µm × 2 cm trap column (Thermo Fisher), followed by separation on a 75 µm × 15 cm Aurora Elite C18 analytical column (IonOpticks). The mobile phases consisted of 0.1% formic acid in water (Solvent A) and 0.1% formic acid in 80% acetonitrile (Solvent B). The gradient began at 1% B, increased to 5% in 0.1 min, then to 30% in 12.1 min, 40% in 0.7 min, and 99% in 0.1 min, which was held for 2 min before re-equilibration, resulting in a total run time of 15 min. The mass spectrometer was operated in data-independent acquisition (DIA) mode. MS1 scans were acquired in the Orbitrap at a resolution of 240,000, with a maximum injection time of 5 ms and an m/z range of 380–980. MS2 scans were collected in the Astral analyzer using a variable windowing scheme (4 Da from 380–750 m/z and 6 Da from 750–980 m/z), with a 6 ms injection time, 28% HCD collision energy, AGC target of 500%, and fragment ion detection from 150–2000 m/z. The total cycle time was 0.6 s. Raw files were analyzed using DIA-NN (v1.9.2) in library-free mode, using the *Escherichia coli* UniProt reference proteome (UP000000625_83333) with deep learning–based spectral and RT prediction enabled^76^. DIA-NN settings included one missed cleavage, a single variable modification (oxidation of methionine), peptide length range of 7–30, precursor charge range of 2–4, precursor m/z range of 380–980, and fragment m/z range of 150–2,000. Quantification was performed in “Robust LC (high precision)” mode with RT-dependent normalization, match-between-runs (MBR) enabled, and protein inference set to “Genes” (heuristic inference disabled). MS1/MS2 tolerances and scan windows were automatically determined by the software. Precursors were filtered at a 1% library precursor q-value, 1% protein group q-value, and 50% posterior error probability. Protein quantification was performed using the MaxLFQ algorithm implemented in the DIA-NN R package, and peptide counts per protein group were determined with the DiannReportGenerator package^76,77^.

### Statistical analysis

Analysis was performed using GraphPad Prism software (GraphPad, La Jolla, CA). In experiments where HUVECs were dosed with BEVs derived from *E. coli* treated with different antibiotics, outlying data points were removed using the ROUT method with Q = 5. For comparisons between groups with a single independent variable and non-normally distributed data, Kruskal-Wallis tests with Dunn’s test for multiple comparisons were used. For comparisons between groups with a single independent variable and normally distributed data, one-way ANOVA with Dunnett’s test for multiple comparisons was used. For comparisons between groups with two independent variables and normally distributed data, two-way ANOVA with Tukey’s test for multiple comparisons was used.

## Data availability

The majority of relevant data for this study are presented in the manuscript and supporting information. An Excel file containing the ICAM-1 fluorescence intensity measurements, nanoparticle tracking analysis concentrations and particle sizes, and Western Blot analysis is available on Figshare along with TSV files with the mass spectrometry data (DOI: 10.6084/m9.figshare.c.7808549).

## Acknowledgements

The authors gratefully acknowledge Dr. Kwang Sik Kim (Johns Hopkins Children’s Center) for providing the *E. coli* K1 RS218 strain. Transmission electron microscopy was performed by the staff of the Electron Microscopy Resource, part of the Center for Advanced Research Technology at the University of Rochester Medical Center.

## Author contributions

LPW, PT, LVM, and TRG Conceptualization; LPW, PT, ACW, and SG Formal Analysis; LVM and TRG Funding Acquisition; LPW, PT, ACW, and SG Investigation; LPW, PT, LVM, and TRG Methodology; LPW, PT, LVM, and TRG Project Administration; SG, LVM, and TRG Resources; LVM and TRG Supervision; LPW, PT, and ACW Validation; LPW, PT, and ACW Visualization, LPW, PT, and ACW Writing – Original Draft Preparation.

## Funding

This work was supported by the National Institute of Allergy and Infectious Diseases (NIAID) of the National Institutes of Health under Award No. R21AI163782 to LVM and TRG, the National Heart, Lung, and Blood Institute (NHLBI) through grant R33HL154249 to TRG, and the National Institute of General Medical Sciences (NIGMS) through grant R35GM153461 to TRG. Additional support was provided by the Hank and Lynn Hopeman Foundation, also awarded to TRG. These funding sources did not influence the study design; collection, analysis, and interpretation of data; writing of the paper; or decision to submit for publication.

## Competing interest statement

The authors declare no competing interests.

## Supporting Information

**Fig. S1:**
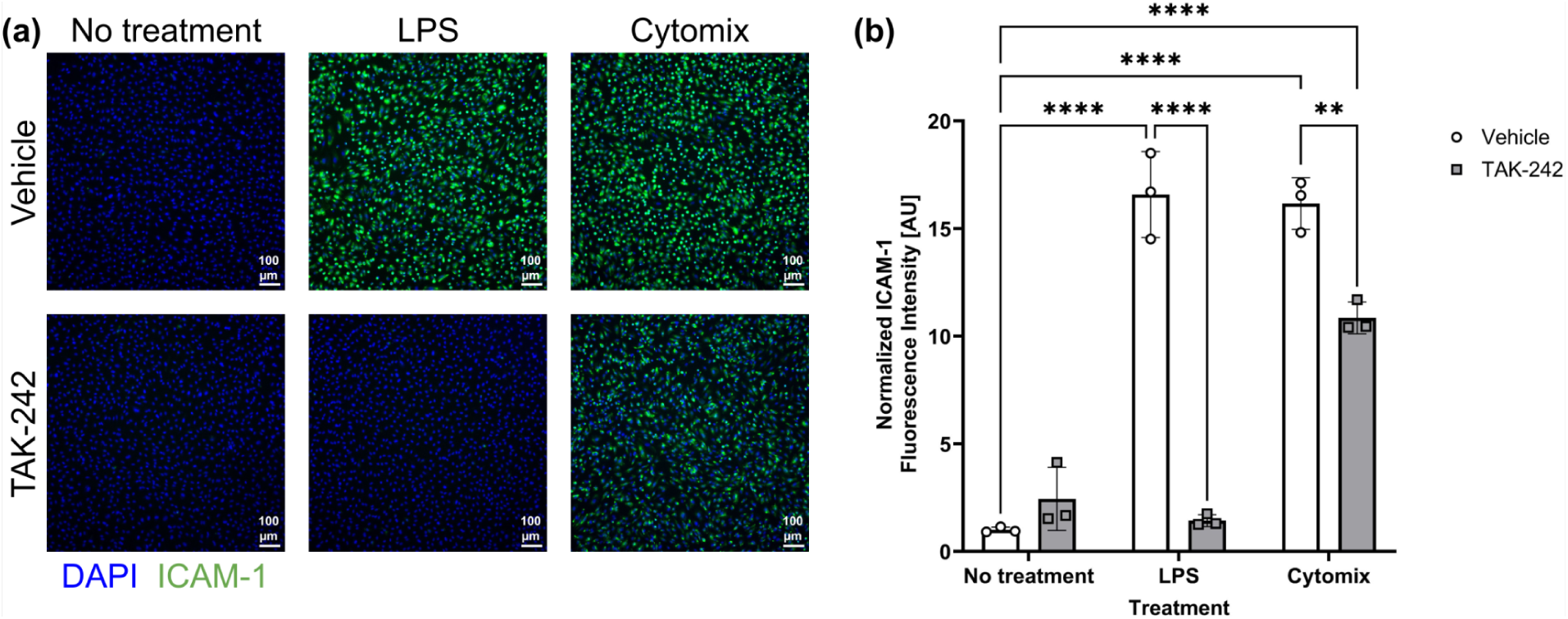
Pro-inflammatory stimulation of non-TLR4 receptors results in elevated ICAM-1 expression even in the presence of TAK-242. (**a**) Immunofluorescence images of HUVECs stained with DAPI (blue) and anti-ICAM-1 antibody (green). Cells were pre-treated with 10 µM of TLR4 inhibitor TAK-242 or a vehicle control (0.04% DMSO) for 4–5 h. They were subsequently exposed to 10 ng/ml LPS or 500 pg/ml cytomix (an equal parts mixture of the pro-inflammatory cytokines TNF-α, IL-1β, and IFN-γ) with either the TAK-242 or vehicle control for 16–17 h. (**b**) Barplot of ICAM-1 fluorescence intensity. The average fluorescence intensity of ICAM-1 in each image was background-subtracted and then divided by the number of cell nuclei in the corresponding DAPI channel. These values were then normalized to the average for the wells that received the vehicle control with no pro-inflammatory treatment. Error bars = s.d. Performed two-way ANOVA with Tukey post-hoc test: ✱✱ = *p* < 0.01, ✱✱✱✱ = *p* < 0.0001

**Fig. S2:**
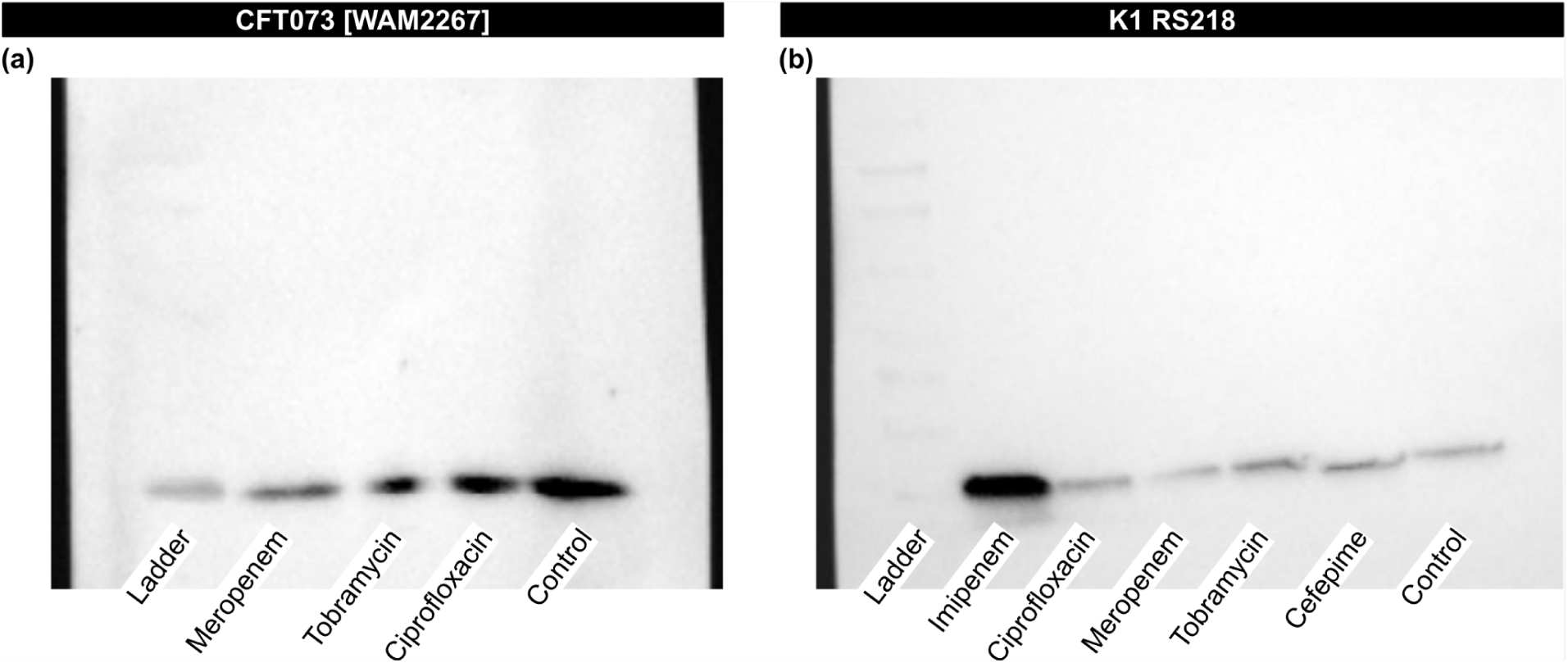
Unedited Western Blot Images. (**a**) Western Blot of Pal in equal volume BEV samples derived from *E. coli* strain CFT073 [WAM2267]. (**b**) Western Blot of Pal in equal volume BEV samples derived from *E. coli* strain K1 RS218. Note that the additional antibiotics imipenem and cefepime were included during early testing prior to the final selection of meropenem, tobramycin, and ciprofloxacin.

## Notes

https://doi.org/10.6084/m9.figshare.c.7808549

### Competing Interest Statement

The authors have declared no competing interest.

### Summary of Updates

Figure formatting has been updated. Some alternate TEM images have been selected for figure 3. Clarifications for original findings have been added in the main text. Additional details have been added regarding the number of bacterial extracellular vesicles available per human umbilical vein endothelial cell. The BEV isolation and western blot protocols have had minor updates.

