## Supporting Information for "Antibiotic treatment modulates *Escherichia coli*-derived bacterial extracellular vesicle (BEV) production and their capacity to upregulate ICAM-1 in human endothelial cells"

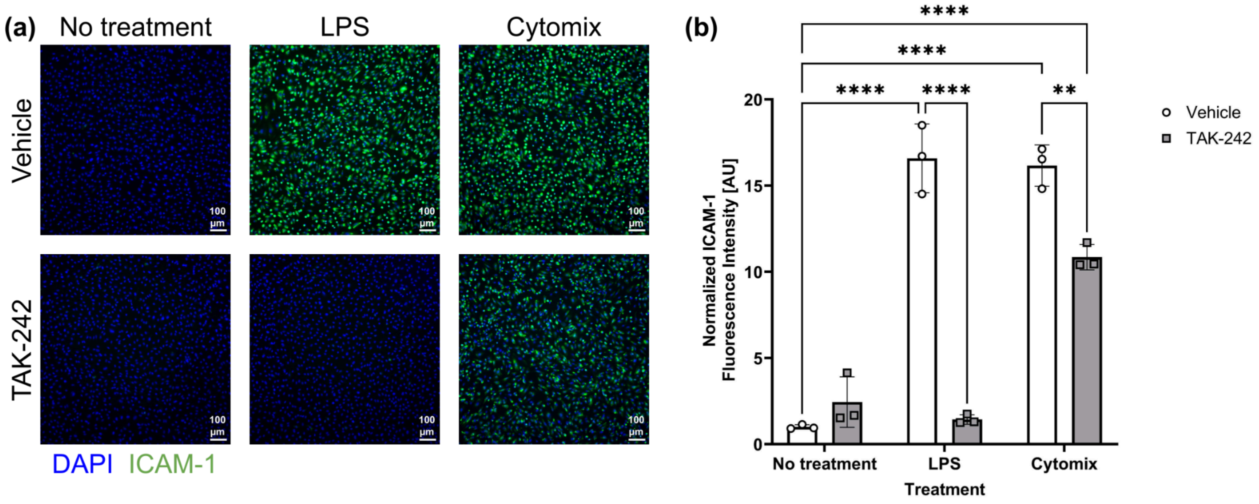


**Fig. S1: Pro-inflammatory stimulation of non-TLR4 receptors results in elevated ICAM-1 expression even in the presence of TAK-242.** (**a**) Immunofluorescence images of HUVECs stained with DAPI (blue) and anti-ICAM-1 antibody (green). Cells were pre-treated with 10 µM of TLR4 inhibitor TAK-242 or a vehicle control (0.04% DMSO) for 4–5 h. They were subsequently exposed to 10 ng/ml LPS or 500 pg/ml cytomix (an equal parts mixture of the pro-inflammatory cytokines TNF-α, IL-1β, and IFN-γ) with either the TAK-242 or vehicle control for 16–17 h. (**b**) Barplot of ICAM-1 fluorescence intensity. The average fluorescence intensity of ICAM-1 in each image was background-subtracted and then divided by the number of cell nuclei in the corresponding DAPI channel. These values were then normalized to the average for the wells that received the vehicle control with no pro-inflammatory treatment. Error bars = s.d. Performed two-way ANOVA with Tukey post-hoc test: ✱✱ = $p < 0.01$, ✱✱✱✱ = $p < 0.0001$

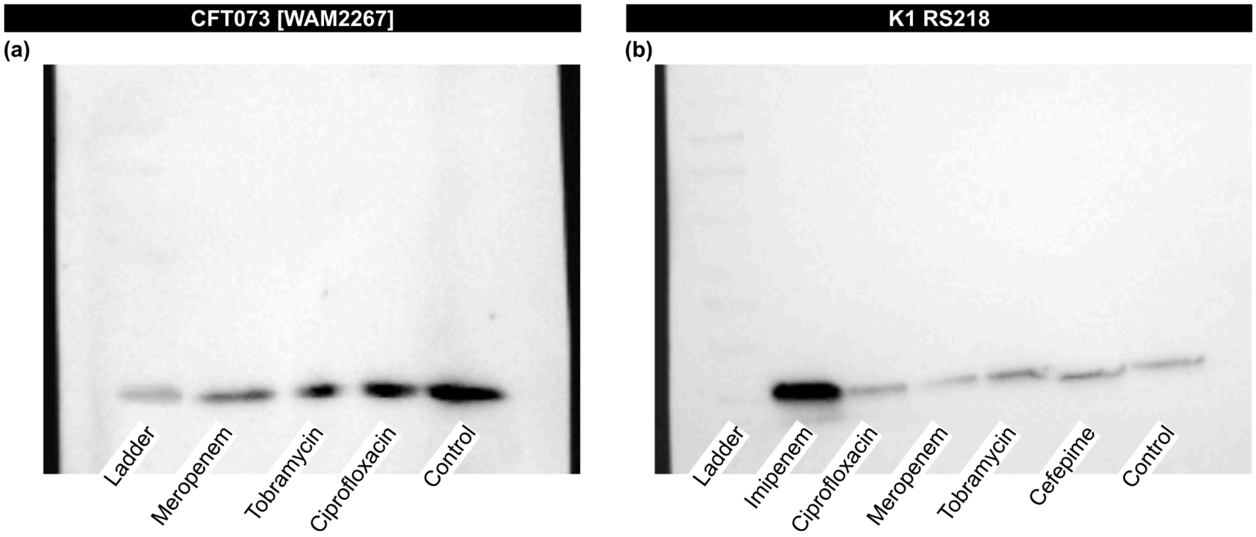


**Fig. S2: Unedited Western Blot Images.** (**a**) Western Blot of Pal in equal volume BEV samples derived from *E. coli* strain CFT073 [WAM2267]. (**b**) Western Blot of Pal in equal volume BEV samples derived from *E. coli* strain K1 RS218. Note that the additional antibiotics imipenem and cefepime were included during early testing prior to the final selection of meropenem, tobramycin, and ciprofloxacin.
